# A sequence-based, deep learning model accurately predicts RNA splicing branchpoints

**DOI:** 10.1101/185868

**Authors:** Joseph M. Paggi, Gill Bejerano

## Abstract

Experimental detection of RNA splicing branchpoints, the nucleotide serving as the nucleophile in the first catalytic step of splicing, is difficult. To date, annotations exist for only 16-21% of 3’ splice sites in the human genome and even these limited annotations have been shown to be plagued by noise. We develop a sequence-only, deep learning based branchpoint predictor, LaBranchoR, which we conclude predicts a correct branchpoint for over 90% of 3’ splice sites genome-wide. Our predicted branchpoints show large agreement with trends observed in the raw data, but analysis of conservation signatures and overlap with pathogenic variants reveal that our predicted branchpoints are generally more reliable than the raw data itself. We use our predicted branchpoints to identify a sequence element upstream of branchpoints consistent with extended U2 snRNA base pairing, show an association between weak branchpoints and alternative splicing, and explore the effects of variants on branchpoints.

## Introduction

Following transcription, which produces RNA molecules identical to the DNA sequence, vast stretches of RNA, called introns, are ‘spliced out’ leaving a string of ‘exons’, which comprise the final messenger RNA. Splicing involves three mechanistically essential sites: the 5’ and 3’ splice sites (5’ss and 3’ss), which define the up and downstream end of an intron, respectively, and a branchpoint, which serves as the nucleophile in the first catalytic step of splicing (Fig. 1a) and is generally located 18 to 45 nucleotides upstream of the 3’ss (Fig. 1g). The branchpoint is recognized by base pairing of the surrounding nucleotides to U2 snRNA and selection on the branchpoint nucleotide itself through a poorly understood mechanism^1^. Overall 3’ss recognition is facilitated by a combination of this selection on the branchpoint, *U2AF65* binding the poly-pyrimidine tract (PPT), *U2AF35* recognizing the core 3’ss signal (Fig. 1a), and a diverse cast of supporting factors^2^.

**Figure 1.**
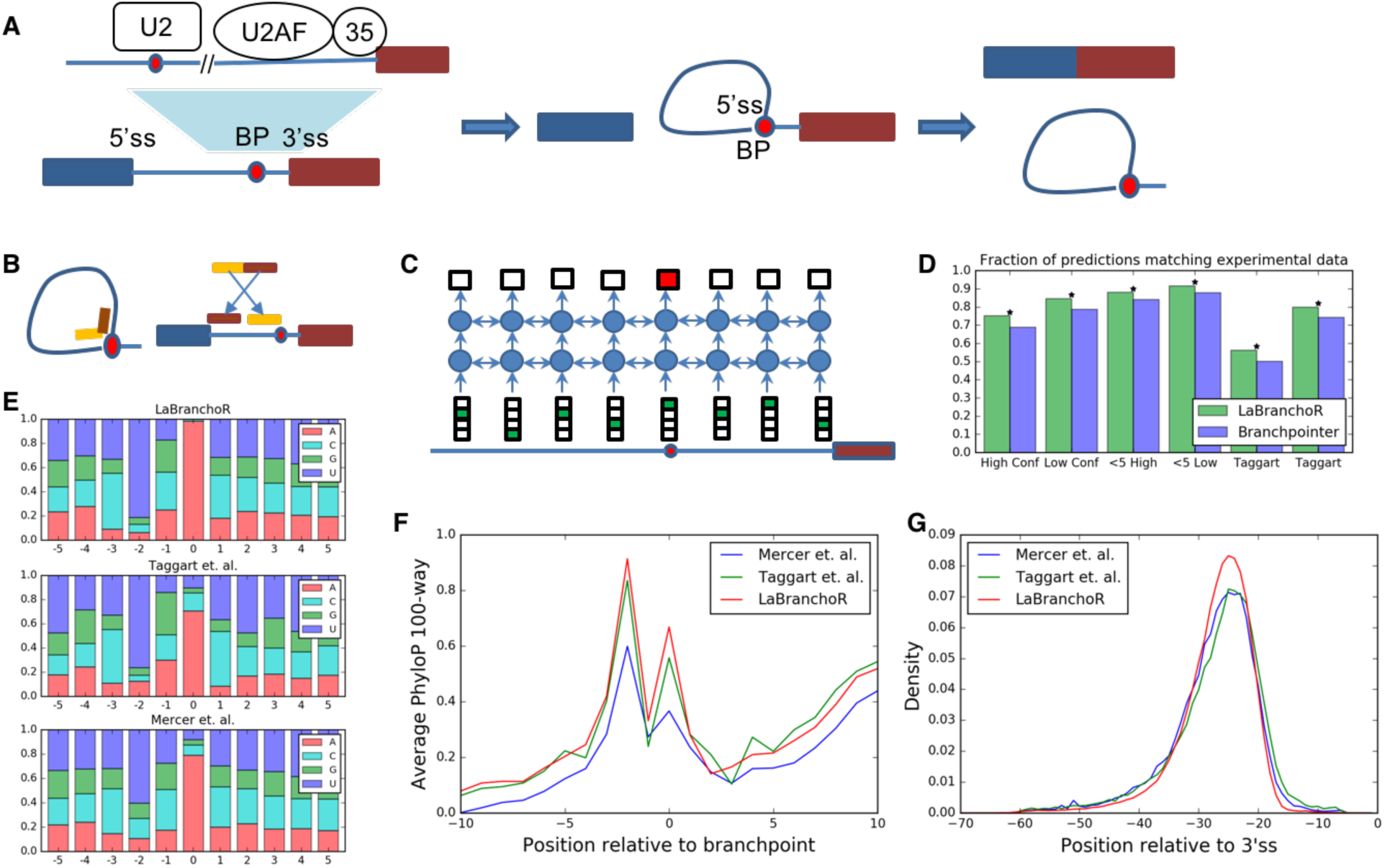
Overview of branchpoints and their genome-wide prediction using LaBranchoR. (A) Overview of the role of branchpoints in splicing and the factors involved in 3’ss recognition. (B) 5’ss-branchpoint junction reads implicate branchpoints. (C) Cartoon of information flow in a bidirectional LSTM. (D) Model performance on a held out test set. A * indicates a significant difference with p < 1e-6 by a onesided Fisher Exact test. Local sequence context (E), PhyloP conservation signature (F), and position relative to 3’ss (G) for predicted and experimentally determined branchpoints.

The locations of the 5’ss and 3’ss can be easily recovered from RNA sequencing (RNA-seq) reads spanning between exons. Similarly, RNA-seq reads spanning 5’ss-branchpoint junctions provide the positions of branchpoints^3^ (Fig. 1b). However, the branched intron byproduct is quickly degraded making these reads exceptionally rare. In fact, a study analyzing a massive collection of internally generated and ENCODE RNA-seq data only provided annotations for 16% of the genome^4^ and even when specialized sequencing was employed only 21% of branchpoints were identified^5^. Furthermore, the small number of reads that are generated from 5’ss-branchpoint junctions provide imprecise information about branchpoint location due to nucleotide skipping and transcript switching caused by the unusual 2’ OH linkage present^4^. Together, these factors have caused the characterization of branchpoints to lag far behind that of 5’ and 3’ splice sites.

The lack of branchpoint annotations has slowed our understanding of the basis of 3’ss selection and diseases caused by mutations to branchpoint sequences themselves and the trans-acting factors that recognize them. However, even with our limited knowledge, it has been shown that branchpoints play a role in Mendelian disease^5,6^, as well as more complex diseases, such as splicing factor 3b associated cancers^7^ and recent reports that expression levels of splicing factor 1 play a vital role in aging^8^.

In response to the lack of experimental methods to effectively provide genome-wide branchpoint annotations and the importance of knowing branchpoints for understanding the basis of 3’ss selection and branchpoint related diseases, we developed a computational method to predict branchpoints, LaBranchoR (**L**ong short-term memory network **Branch**point **R**etriever). Specifically, we focus on the problem of predicting the most likely branchpoint given the associated 3’ss, which we took to be the most salient task given the widespread availability of 3’ss positions. LaBranchoR is based on a bidirectional long short-term memory network (LSTM), a ‘deep learning’ algorithm shown to be wildly successful in modeling sequential data such as time-series and natural language^9,10^. The use of a LSTM allowed us to build a model based on solely the RNA sequence, free from the biases of hand-engineered features.

Throughout our study, we compare LaBranchoR to two recently proposed computational methods, which focus on two distinct tasks: a machine learning approach for branchpoint prediction, *branchpointer*^6^, and a method to remove noise in the experimental data proposed by Taggart et. al., 2017^4^. *Branchpointer* employs an ensemble of support vector machines and gradient boosting tree classifiers, which take as input a library of engineered features. *Branchpointer* is trained on the same set of experimental branchpoints as our model: the ‘high confidence’ set of branchpoints reported by *Mercer et. al*. 2015. *Taggart et. al*. explicitly modeled U2 snRNA base pairing potential, ignoring the branchpoint position itself, and the likelihood of observing a given nucleotide skipping distance to resolve noise introduced by nucleotide skipping. They used this model to produce maximum likelihood branchpoint predictions for an independent set of RNA-seq data, including a diverse range of cell lines.

In this study, we show that LaBranchoR has strong predictive performance, exceeding that of previous methods. It also disregards noise in the experimental data leading to robust predictions that are often more accurate than the raw data itself. After showing the accuracy of our predictions, we use them to evaluate genome-wide properties of branchpoints and find that we recover known trends, as well as several novel insights. We conclude that branchpoint strength plays a role, similar to that of 3’ss strength, in alternative splicing. We identify a novel upstream recognition element, which is consistent with a recent cryo-EM model of the spliceosome depicting the relevant bases in duplex with U2 snRNA^11^. Finally, we show that LaBranchoR predictions overlap with more pathogenic variants than previous computational predictions, as well as the raw data itself.

## Results

We used a bidirectional LSTM network to learn a mapping between RNA sequence and branchpoint locations (Fig. 1c). We trained our model on the high confidence set of branchpoints annotated by *Mercer et. al*. 2015. The RNA sequence 1 to 70 base pairs upstream of each 3’ss was encoded as a ‘one-hot’, 70 x 4-dimensional matrix and served as the sole input to our model. All branchpoints 5 to 60 base pairs upstream of a 3’ss were encoded into a length 70 binary vector. We reserved chromosome 1 for testing, chromosomes 2, 3, 4 for model selection and the remaining data were used for model training. While our model predicts a length 70 vector of branchpoint probabilities for each 3’ss, we chose to focus our analysis on a single predicted branchpoint per 3’ss, corresponding to the argument maximum of the predictions.

### LaBranchoR provides accurate genome-wide branchpoint annotations

We found that our model’s predictions generally agreed with branchpoints implicated by both *Mercer et. al*. 2015 and *Taggart et. al*. 2017. On a test set held-out from model training and parameter tuning, we found that our predicted branchpoint coincided with a high confidence Mercer site for 75% of 3’ss (Fig. 1d). Expanding this analysis to consider low confidence sites and cases where we predict within 4 nucleotides of a Mercer site yields an accuracy of 84% and 91%, respectively. Our predictions have a lower agreement with the Taggart set, where we find that we overlap an experimental site for only 56% of 3’ss and 80% of the time we are within 4 nucleotides. Notably, restricting to Taggart sites with an A at the branchpoint yields a stronger overlap of 71% and 84%, respectively (Supplementary Fig. 1a).

These performances represent a 5 to 12 percentage point improvement over the current state-of-the-art branchpoint prediction software, *branchpointer*^6^ (Fig. 1d). Comparing to them on our test set yielded a 7 percentage point advantage and even when only considering branchpoints in positions −18 to −45 from the 3’ss, where *branchpointer* makes predictions, we maintain a 5 percentage point increase in performance. However, this evaluation of their performance is overly-optimistic, as *branchpointer* had seen roughly 80% of the data from our test set in its training. On the intersection of our test sets, LaBranchoR outperforms *branchpointer* by a 7 and 12 percentage point margin for the −18 to −45 and −5 to −60 ranges, respectively (Supplementary Fig. 2b). Area under receiver-operator curve and precision-recall curve statistics for all mentioned evaluations are in Supplementary Figure 2c.

We found that the bulk trends in sequence motifs, positional distribution, and conservation signatures are similar, but display a few key differences between our predicted, Mercer, and Taggart branchpoints (Fig. 1e-g). Our predictions and the Taggart set have a similar sequence motif with stronger nucleotide content biases than the Mercer set, which closely match the motif expected for base pairing with U2 snRNA (Fig. 1e). However, our predictions show a higher rate of A branchpoint nucleotides. While this could represent a “modal collapse” of our model or bias against C branchpoints in our training data, a strong bias to an A at the branchpoint is supported by a study of positional k-mer enrichment^12^. Additionally, the branchpoint nucleotide was not considered in the Taggart et. al. denoising strategy, so there could exist cases where a slightly stronger U2 base pairing sequence without an A was incorrectly selected over a different position with an A nucleotide at the branchpoint.

Branchpoints have previously been observed to have a distinct conservation signature, closely aligning with the bias in nucleotide content at each position^5^. Since conservation was not seen by LaBranchoR during training, it can be used as an independent validation metric. We found that our predicted branchpoints have the strongest PhastCons and PhyloP (Supplemental Fig. 2a and Fig. 1f) conservation signature, followed by the Taggart sites.

### LaBranchoR disregards noise in the experimental data leading to robust predictions

Considering these observations, we performed a more fine-grained analysis of the conservation signatures and sequence motifs where we agree and disagree with the Mercer and Taggart set for 3’ss in our test set (and validation set for the Taggart set to arrive at roughly equal numbers). For both the Taggart and Mercer set, the strongest conservation signatures are present where predictions match the experimental data (Fig. 2c-d). Interestingly, the intersection of LaBranchoR and Taggart sites results in the strongest conservation signature and a −2 U and branchpoint A are nearly always present (Fig. 2c and Supplementary Fig. 1b). We found that predictions matching a Mercer low confidence branchpoint resulted in only a slightly weaker conservation signature than those matching high confidence sites (Fig. 2c).

**Figure 2.**
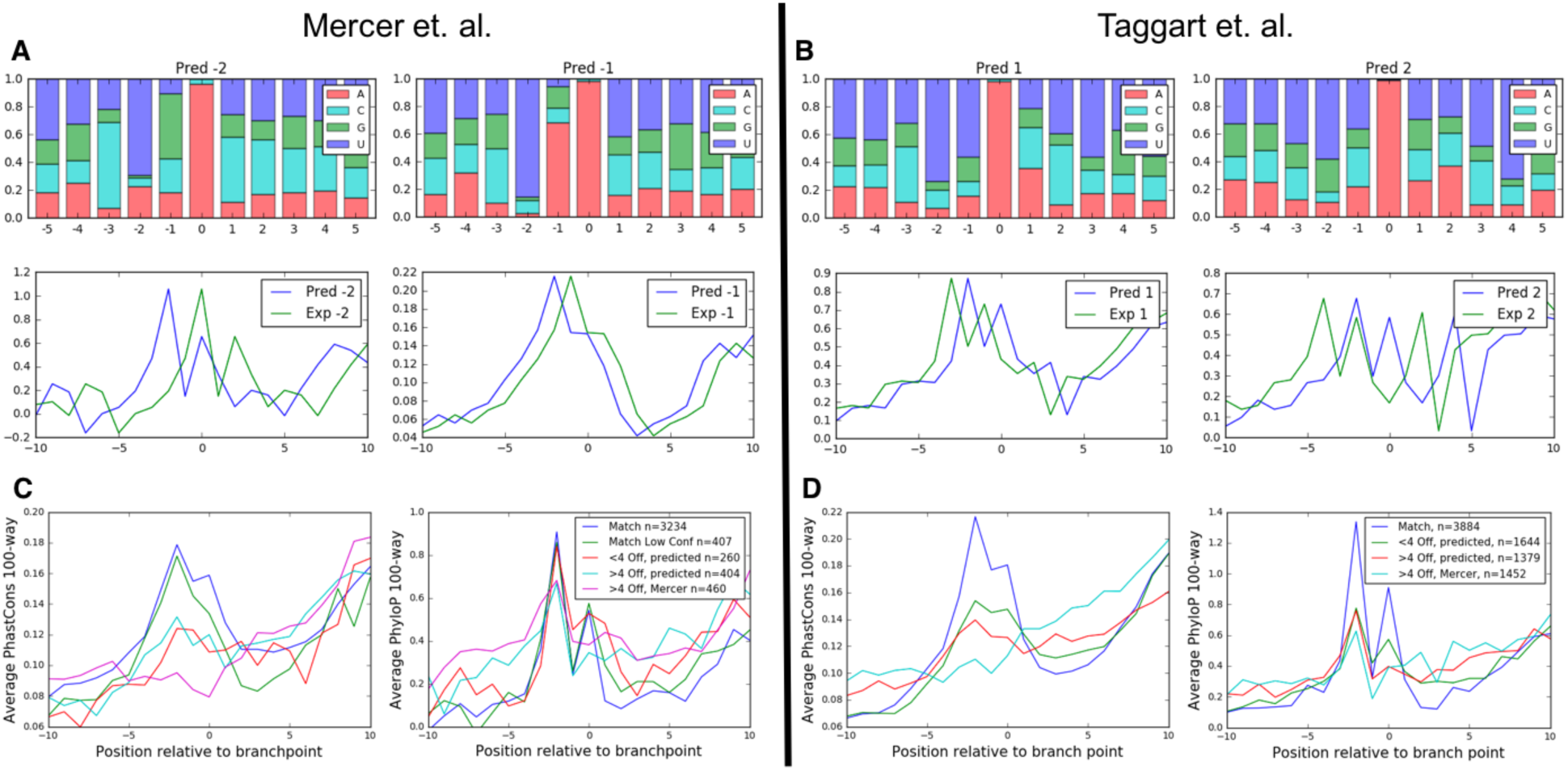
LaBranchoR disregards noise by applying robust genome-wide patterns. (A) Sequence motifs and PhyloP conservation signatures corresponding to 1 and 2 nucleotide shifts further into the intron in the Mercer et. al. high confidence branchpoints and (B) 1 and 2 nucleotide shifts towards the 3’ss in the Taggart et. al. data. (C, D) PhastCons and PhyloP conservation signatures for cases where our prediction agrees with the Mercer (C) and Taggart (D) data, is within 4 nucleotides, and is off by more than 4 nucleotides. Where our prediction and the experimental data diverge, a signature indexed by both our predictions and the experimental coordinate is presented.

The next strongest conservation signatures were present for cases where LaBranchoR diverges from a Mercer or Taggart site by a small shift, which we defined as 4 nucleotides or less. For both experimental sets, we found that indexing by our predicted branchpoint results in a stronger conservation signature than indexing by the experimental coordinate (Fig. 2c-d). The Mercer sites were enriched for small shifts further into the intron, representing nucleotide skipping, whereas the Taggart sites were enriched for shifts towards the 3’ss, perhaps representing over-compensation for nucleotide skipping. The correctness of our predictions in these cases is supported by the sequence motifs and conservation signatures present (Fig. 2a-b). In all cases, when indexing by our predicted branchpoint, a strong branchpoint A and −2 U signature is present, but this same trend did not hold when indexing by the experimental coordinates (Fig. 2a-b). Similarly, the PhyloP and PhastCons signatures more closely resemble the consensus pattern when centered on our predictions as opposed to the experimental coordinates (Fig. 2c-d).

In cases where our predictions disagree by larger shifts (4+) from an experimental branchpoint, we found that our predicted branchpoints display a stronger PhastCons signature. For both the Mercer and Taggart data, the conservation signature centered on the experimental coordinate showed no clear increase in relation to the branchpoint (Fig. 2c-d). However, the story was not as clear for PhyloP conservation scores, as we found that all three sets show an increase at the −2 position, although the signature appears to be more similar to the consensus signature when indexing by our predicted coordinate than the experimental coordinates. Due to this ambiguity, we refrain from making strong statements about the accuracy of our predictions deviating by more than 4 nucleotides from an experimental branchpoint. However, we think that it is safe to conclude that a non-negligible proportion of these predictions are likely correct, implying that our performance estimates are loose lower bounds.

### Cytosine branchpoints and branchpoints without a −2 uracil have distinct properties

The strong trend towards A nucleotides at the branchpoint and U at the −2 position in our predicted set led us to consider if branchpoints lacking these properties displayed any distinct patterns. While C branchpoints represented only 1.5% of predicted branchpoints and 10% of Taggart branchpoints, about a fifth of predicted and Taggart branchpoints lack a U at the −2 position, so this subset represents a significant proportion of branchpoints in the genome. We chose to analyze the properties of these sites in parallel as they both likely represent weaker than average branchpoints and were, in fact, found to follow many of the same trends.

Both sets present sequence and conservation signatures diverging from the bulk trends. Branchpoints lacking a −2 U show an increased rate of U at the −3 position and C at the −1 and −4 positions (Fig. 3a). Meanwhile, C branchpoints have an increased rate of C nucleotides at the −3 and +1 position, consistent with the Taggart C branchpoints (Fig. 3a). For C branchpoints, the conservation signature shifts to form bumps at the positions of these increased nucleotide biases (Fig 3b). For branchpoints lacking a −2 U, the previous bump at the −2 position is entirely lost and, inexplicably, the branchpoint nucleotide also lacks an increase in conservation (Fig. 3b).

**Figure 3.**
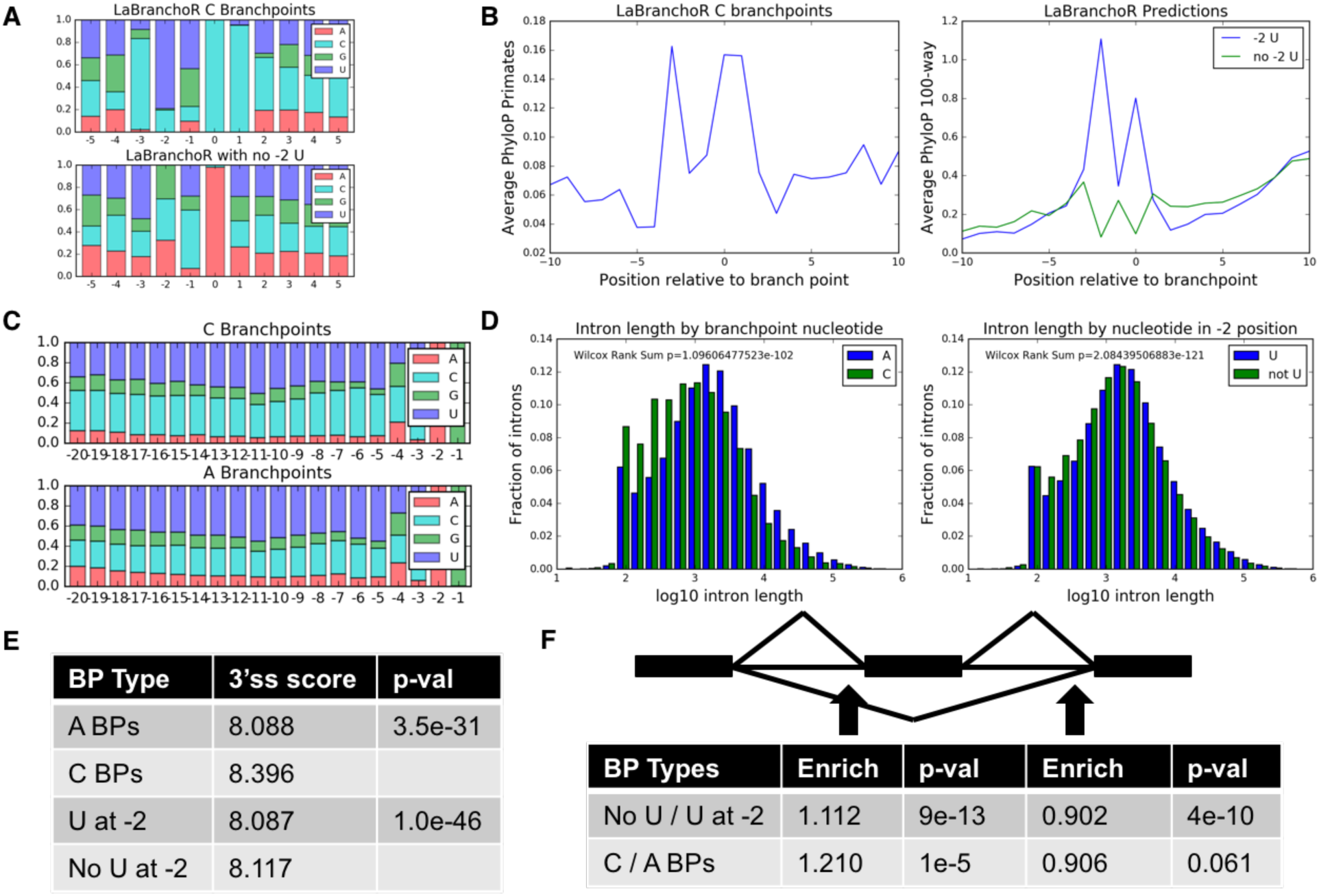
Cytosine branchpoints and branchpoints lacking a −2 uracil display distinct properties. (A) Sequence motifs for C branchpoints (top) and branchpoints lacking a −2 U (bottom). (B) Conservation signatures for C and no −2 U branchpoints. (C) 3’ss nucleotide content for A and C branchpoints. C branchpoints have 1.43 times more C nucleotides in the −20 to −5 range than A branchpoints. (D) Both C and no −2 U branchpoints display significantly shorter introns than expected. (E) Both groups have stronger than expected average 3’ss MaxEntScan scores (two-sided Wilcoxon Rank Sums test). (F) Both groups are enriched upstream of cassette exons and de-enriched in introns downstream of cassette exons (two-sided Fisher Exact Test).

We went on to examine if these sets of branchpoints were enriched for particular types of splicing events. We found that both sets were associated with short introns with median intron lengths shifting from 1654 to 1314 nucleotides for branchpoints with and without a −2 U and from 1603 to 807 nucleotides for A and C branchpoints (Wilcoxon Rank Sum test p < 10^-100 and p < 10^-120) (Fig. 3d). Additionally, we found that both sets were enriched in retained introns and upstream of skipped exons (Fig. 3e and Supplementary Table 1). Interestingly, these sets were de-enriched from introns downstream of skipped exons (enrichment of 0.9 for both, p < 10^-9 for −2 U, no −2 U and p = 0.065 for A, C). The enrichment of weak branchpoints upstream of skipped exons was previously observed in the Mercer branchpoints, albeit at a lower confidence^5^. Interestingly, this same study found that there was no enrichment of weak branchpoints in retained introns, likely due to the small number of branchpoints and occurrences of retained introns in the genome.

Both subsets of branchpoints are associated with strong 3’ss sequences, as determined by MaxEntScan^13^ (Fig. 3f). This trend was previously observed for C branchpoints^4^, however, we add the observation that C branchpoints are associated with strikingly C-rich poly-pyrimidine tracks with a 1.42 fold enrichment of C’s in positions −20 to −5 from the 3’ss (p ≅ 0 by two-sided Fisher Exact test) (Fig. 3c).

### A nucleotide content signature consistent with extended base pairing to U2 snRNA is present upstream of branchpoints

Analysis of the sequence upstream of our predicted and experimental branchpoints revealed peaks in G content centered at positions −6 to −7 and −12 and a peak in C content at position −9 (Fig. 4a-c and Supplementary Fig. 3a-c). To the best of our knowledge, this sequence motif has not been previously observed in association with branchpoints and perhaps represents a novel sequence feature aiding in branchpoint recognition. A recent cryo-EM spliceosome structure shows these bases in duplex with U2 snRNA^11^ (Fig. 4e). There is a guanine at positions −12 and −7 and cytosine at −9 of U2 snRNA in position for Watson-Crick base pairing to these peaks (Fig. 4d). Interestingly, the cryo-EM structure shows a distorted helix between the canonical branchpoint recognition sequence and this region of extended base pairing. This distortion could allow for shifts in the alignment of the intronic RNA and U2 snRNA resulting in the smooth observed peaks.

**Figure 4.**
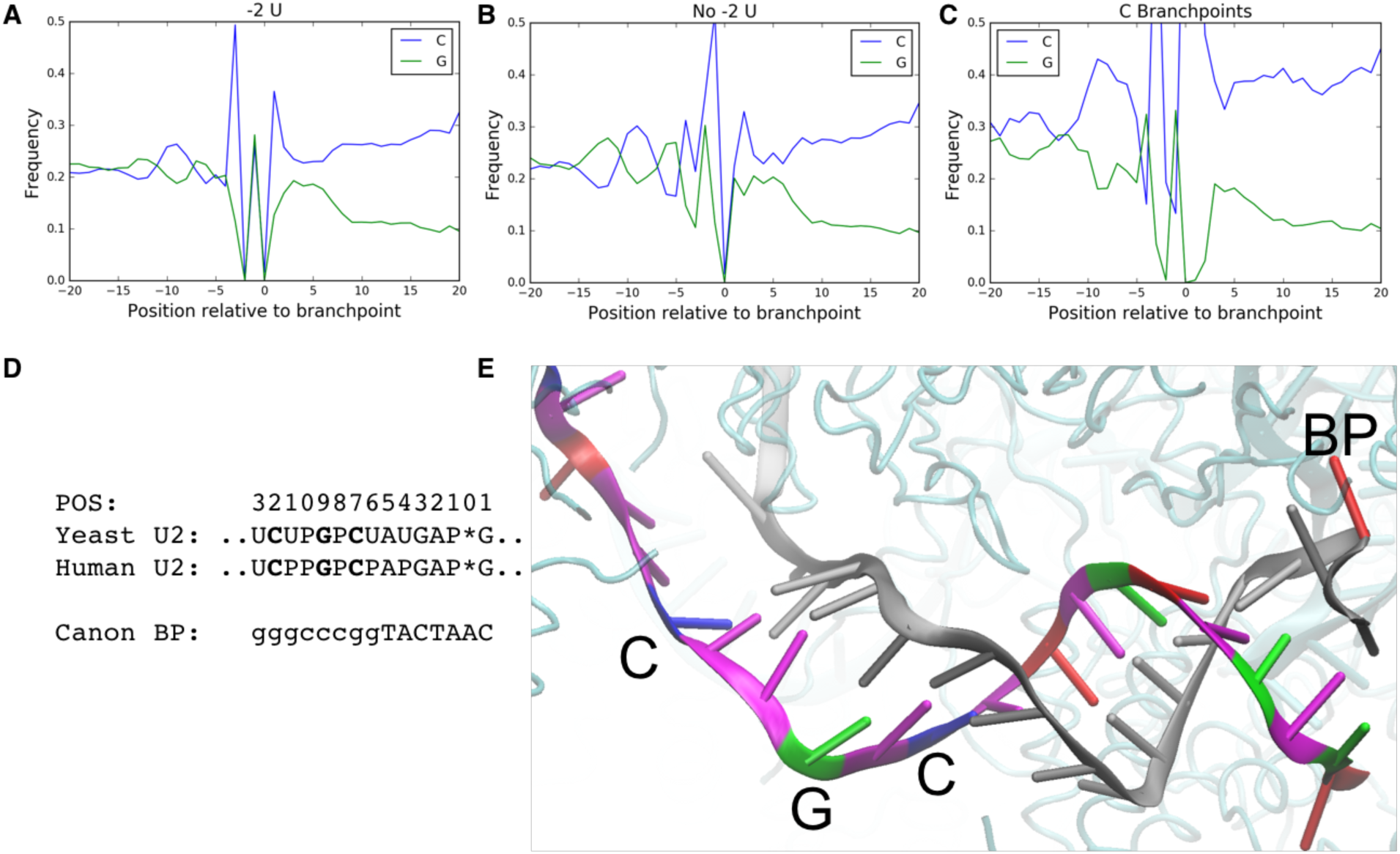
Branchpoints display a nucleotide content signature, consistent with extended base pairing with U2 snRNA. This signature is stronger in branchpoints lacking a −2 U (B) than in branchpoints with a −2 U (A). (C) C branchpoints display a strong increase in C nucleotide content in this same position. (D) The human and yeast U2 snRNA sequence positioned for interacting with this upstream recognition element (top). P represents psuedouridines and * corresponds to the branchpoint position. The canonical branchpoint motif (upper case) and positions of G, C content signature (lower case). (E) A image from a cryo-EM structure of the spliceosome (PDB 5mfq) shows an extended intron-U2 snRNA duplex. The intronic sequence is shown in gray, except for the branchpoint which is shown in red. The U2 snRNA sequence is colored by base (red, blue, green and magenta for A, C, G and T, respectively).

This signature is significantly stronger for branchpoints lacking a −2 U than for branchpoints with a −2 U present with a 1.190, 1.145, 1.192 fold increase in the strength of peaks at −6, −9 and −12, respectively (Fisher Exact 2-tailed P < 10^-54, 10^-40, 10^-59). This trend is stronger still in C branchpoints as opposed to A branchpoints, although in this case the increase in C content seems to dominate the G content signature and the upstream sequence is overall more C-rich making enrichment analysis challenging (Fig. 4c).

### Branchpoints are enriched for pathogenic variants, whereas likely benign variants are excluded

We assembled a set of pathogenic variants by taking the union of variants labeled ‘Pathogenic’ in ClinVar and ‘DM’ in HGMD and filtering out any variant with a nonsynonymous effect on a protein coding sequence^14,15^. We found that LaBranchoR predictions display a strong overlap with these sites with 52 variants directly overlapping the branchpoint, 15 at the −2 position, and 25 in positions −1, −3, and +1 for a total of 92 pathogenic variants (Fig. 5c). In comparison, despite predicting 69,617 (133%) more branchpoints (as they allowed for multiple branchpoints per 3’ss), *branchpointer* predictions have only 46 variants directly overlapping the branchpoint, 10 at the −2 position, and 27 in positions −1, −3, and +1 for a total of 88 pathogenic variants. Tuning LaBranchoR to predict the same number of branchpoints resulted in the prediction of 106 total pathogenic variants in the −3 to +1 interval (Despite this observation, we stuck with predicting one branchpoint per 3’ss as our primary task because the rate of overlap with pathogenic variants is much lower in the additional branchpoints (1 in 4,273) than the initial predictions (1 in 2,174)). Additionally, when considering regions upstream of 3’ss where Mercer and Taggart branchpoints exist, our predictions overlap a larger number of pathogenic variants than the experimental data (Fig. 5a-b).

**Figure 5.**
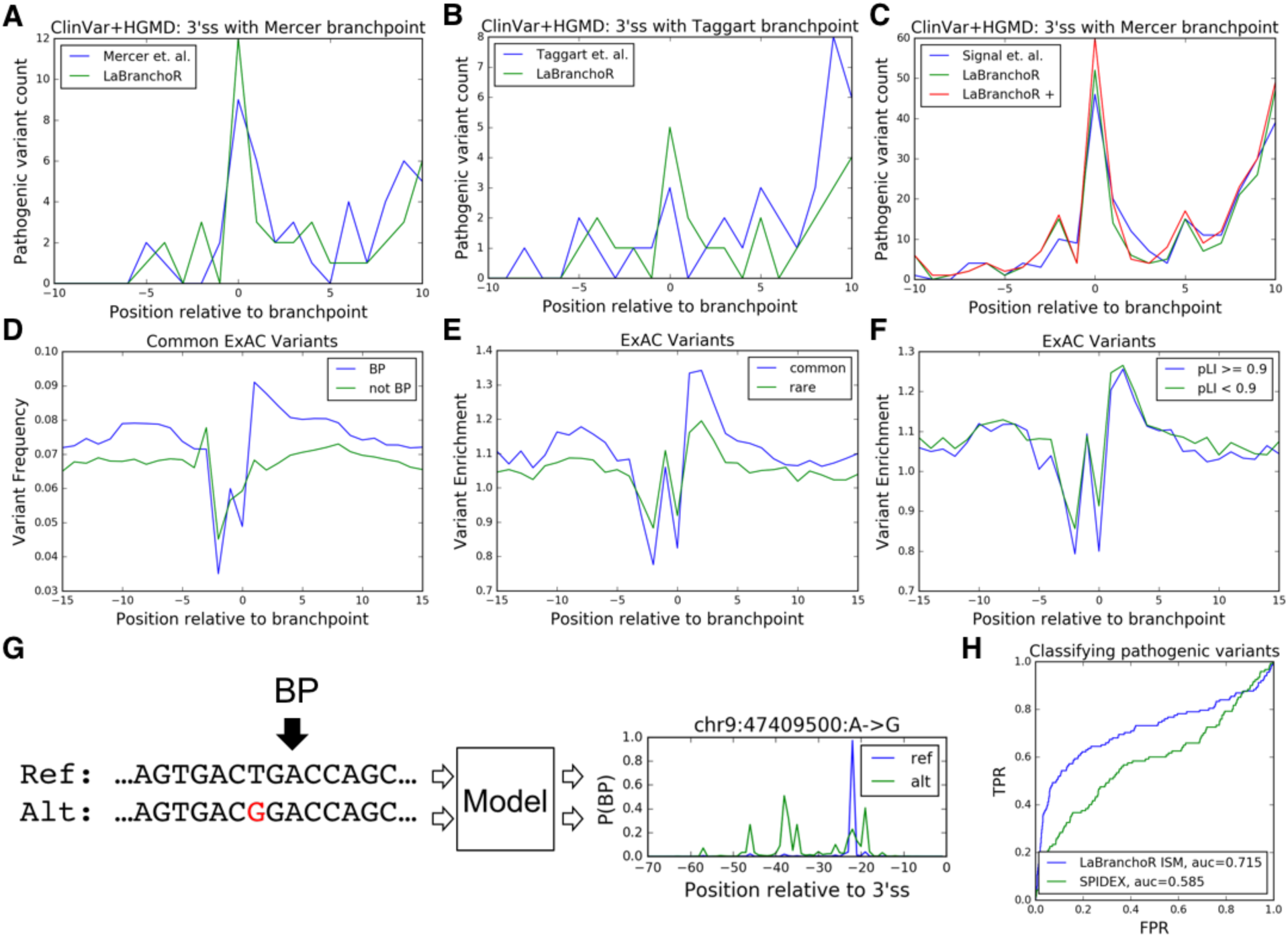
LaBranchoR predicted branchpoints overlap pathogenic variants and exclude common variants. (A) Overlaps with pathogenic variants for Mercer high confidence branchpoints and LabBranchoR predictions for 3’ss with a Mercer high confidence site. (B) Similar for Taggart branchpoints. (C) Genome-wide pathogenic variant overlaps for the current state-of-the art branchpoint predictor, *branchpointer*, LaBranchor, and LaBranchor tuned to predict the same number of branchpoints as *branchpointer* (LaBranchoR +). (D) Comparison of common variant rate in ExAC for distance from 3’ss matched UNA trinucleotides, where the A is implicated as a branchpoint (BP) and not a branchpoint (not BP). (E) Enrichments in variation rate of branchpoint UNAs, as compared to non-branchpoint UNAs for common and rare variants. (F) Similar comparing branchpoints in high probability loss of function intolerant (pLI) genes to low pLI. (G) *In silico* mutagenesis scores are defined as the change in score of our predicted branchpoint induced by the variant. (H) LaBranchoR *in silico* mutagenesis (ISM) scores effectively classify pathogenic variants. A receiver-operator curve for HGMD and ClinVar variants sorted by LaBranchoR ISM scores and SPIDEX scores.

Conversely, we reaffirm that variants present in the general population, as reported by the ExAC consortium^16^, are excluded from branchpoints^5,6^. To circumvent sequence and distance biases in variation rate, we compared the variation rate at predicted branchpoints with sequence UNA to UNA tri-nucleotides not implicated as branchpoints from a matched distance distribution. We found that branchpoints show a 0.776 and 0.815 fold enrichment of common variants (occurring in at least one in 10,000 people) at the −2 and branchpoint positions (p < 10^-40, 10^-33 by two-sided Fisher Exact Test) (Fig. 5d). Furthermore, these trends are stronger for common variants than for rare variants and for branchpoints in genes with probability of loss of function intolerance (pLI) >= 0.9 than genes with pLI < 0.9 (p < 10^-5, 10^-6 by two-sided Fisher Exact Test) (Fig. 5e-f)^16^. Interestingly, we found that for all sets there is a marked enrichment in variation in the +1 to +4 positions, which is mirrored by a lull in pathogenic variants, perhaps indicating that this region generally plays little functional role apart from serving as a linker between the PPT and branchpoint sequence (Fig. 5a-f).

We quantified the effect of variants on branchpoint strength by comparing predictions for the reference and alternative sequences: a technique often referred to as *in silico* mutagenesis (ISM)^6^. Specifically, we calculated the change induced by the variant on the score of the branchpoint predicted for the reference sequence (Fig. 5g). We found that pathogenic variants located 18 to 45 nucleotides upstream of a 3’ss have significantly lower mutagenesis scores than likely benign, ExAC variants (Fig. 5h). In fact, our ISM scores outperform a state-of-the-art model for predicting changes in splicing induced by variants, SPIDEX^17^, in separating pathogenic variants from ExAC variants in the −18 to −45 nucleotide range, achieving an area under the receiver-operator curve statistic of 0.718, as compared to 0.585 for SPIDEX (Fig. 5h).

## Discussion

While it is hard to precisely evaluate our model’s performance due to pervasive noise in the experimental data, our analysis suggests that LaBranchoR is able to correctly predict a branchpoint for over 90% of 3’ss. We arrive at this conclusion based on explicit agreement with experimental annotations (75% high confidence, 84% low confidence), analysis of conservation signatures in cases where we deviate from experimental annotations by less than 5 nucleotides (91%), and slightly less compelling conservation signatures in the remaining case. Furthermore, it appears that having a reliable way to remove noise introduced by nucleotide skipping and transcript switching will continue to be valuable even as more experimental data becomes available.

It has been previously shown that 3’ss strength correlates with alternative splicing outcomes^18^ and our analysis shows that the same trends hold for branchpoint strength. Our genome-wide branchpoint predictions allowed us to assess the properties of two groups of weak branchpoints: those lacking a −2 U and those with a C at the branchpoint. We found that these weak branchpoints are enriched for two types of conditionally used splice sites: those involved in intron retention and upstream of skipped exons. Additionally, we found that weak branchpoints are excluded from introns downstream of skipped exons, supporting that branchpoint strength helps enable competition between the two relevant 3’ss. This complements a report that branchpoints abnormally far upstream of 3’ss enable exon skipping by slowing the upstream splicing reaction between the first and second catalytic steps^4^. We additionally observe that weak branchpoints are generally associated with stronger than average splice sites, supporting that 3’ss selection is a holistic process where the strength of the branchpoint, PPT and core signal interact to result in the overall strength of the 3’ss.

We found a distinct signature in G and C nucleotide content upstream of branchpoints, consistent with an extended region of base pairing with U2 snRNA. This extended base pairing is observed in a recent cryo-EM structure of the human spliceosome^11^ and is consistent with an early biochemical study showing that “SAP 145, together with four other SF3a/SF3b subunits, UV cross-links to pre-mRNA in a 20-nucleotide region upstream of the BPS”^19^. This region of U2 snRNA shares 100% sequence identity with U2 snRNA in budding yeast, albeit in humans 4 of these bases are modified to form psuedouridines, while in yeast only 2 have this modification (Fig. 4d)^20^. Indeed, we observed a similar pattern in G, C content in a dataset of 718 budding yeast branchpoints^21^ (Supplementary Fig. 3d). The extensive pseudouridylation, a modification resulting in stronger base pairing to all bases^22^, of this stretch of U2 snRNA could provide a mechanism by which this region is able to interact favorably with a diverse set of RNA sequences.

Mirroring the trends in 3’ss strength, we found that this signature was on average stronger for weak branchpoints, supporting that it plays a positive role in branchpoint selection and enables usage of otherwise weak branchpoints. Together with the biochemical data showing that SF3b contacts this region, disruption of this extended interaction in *SF3b* mutants presents a potential mechanism of the erroneous splicing in *SF3b* associated cancer. Disruption of the extended interaction could require a stronger core branchpoint for stable U2 binding, resulting in the observed usage of novel 3’ss, characterized by stronger than average branchpoints^7^.

The initial motivating factor for developing LaBranchoR was to aid in the identification of pathogenic genetic variants and we found that LaBranchoR has state-of-the-art performance in this area. While the strong overlap between our predicted branchpoints and variants associated with Mendelian disease is not surprising based on past work^3,5,6^, our predictions overlap pathogenic variants at a higher rate than both previous computational predictions and the raw experimental data. Furthermore, we found that LaBranchoR *in silico* mutagenesis scores are better able to distinguish pathogenic variants from variants in the general population than SPIDEX scores, showing that explicit branchpoint prediction provides information not captured by generic splicing models.

## Online Methods

### Preparation of Annotations

A bed file of high confidence branch points implicated by *Mercer et. al*. (their Supplementary data table 2) was downloaded from Genome Research, as were the *Taggart et. al*. branchpoint predictions. We did not consider Taggart predictions whose “binding model” was ‘none’, ‘transcript_skipping’, or ‘circle’. A set of 718 budding yeast branchpoints were obtained from *Gould et. al. 2016*.

Introns were extracted from the Gencode v19 annotations for all protein coding genes. Bedtools was used to link branch points to three prime splice sites using the intersect-loj command. Branchpoints were considered to be associated to a 3’ss if they lie between 5 and 60 base pairs upstream of it. The Mercer high confidence set of branchpoints was used to produce a training, validation, and test set split by chromosome. Chromosome 1 was used as a test set. Chromosomes 2, 3, and 4 were used as a validation set and all others were used for training.

PhyloP and Phastcons 100-way scores were downloaded from the UCSC website. They were used to produce average conservation plots using in house scripts with the help of bedtools.

### Model training

For each 3’ss, a target vector was composed to have a 1 in each position with a high confidence Mercer et. al. branch point and zeros elsewhere. An input vector was composed by one-hot encoding the 70 base pairs of genomic sequence immediately upstream of the 3’ss.

The model used was a two-layer bidirectional LSTM. The model was implemented using keras version 2.0.4. The final model has 32 hidden nodes in each direction for both layers. The output of both LSTM layers are stacked to form a 70x64 tensor that is passed to the next layer. At the top of the network is a time distributed 1D convolution with sigmoid activation mapping the 64 outputs per position to a single number. A binary cross entropy loss function was employed in training. Both recurrent (0.05) and normal dropout (0.15) were employed. The model was trained with the Adam optimizer with default keras parameters. The model was trained until the number of validation set branch points that overlap with Mercer et al branch points did not increase for 15 epochs.

### Model Testing

Model performance was tested using the 4306 3’ss on chromosome 1 that were held out from the training and validation set. The fraction of the top scoring branchpoints for a given 3’ss overlapping an experimental branch point was reported using an in house script. Sklearn functions were used to compute receiver-operating curve and precision-recall curve statistics. For each of these statistics, we calculated them separately for considering all bases in the −70 to −1 positions that were assigned branch point scores as well as for the −45 to −18 positions in which the vast majority of branch points fall. As was done in *Signal et. al*., we masked positions corresponding to low confidence branchpoints from the negative set, when computing area under curve statistics.

### Comparison to branchpointer

We compared our model to predictions from the *branchpointer* R package created by *Signal et. al*. (https://bioconductor.org/packages/release/bioc/html/branchpointer.html)^6^. Predictions were prepared for our test set by closely following the example given in the reference manual. We additionally downloaded a precomputed file of genome-wide predictions in Gencode v19 introns for analysis of overlap with pathogenic variants. We obtained the training and test set used by *Signal et. al*. from https://osf.io/hrqvq/.

### Variants

A set of pathogenic variants was composed by taking the union of ClinVar ‘Pathogenic’ and HGMD PRO 2017 ‘DM’ variants. We removed all variants that affected a protein coding sequence. ANNOVAR v527 was used to annotate variants with predicted effect on protein-coding genes using gene isoforms from Ensembl gene set version 75 for the hg19/GRCh37 assembly of the human Genome 35. All coding isoforms were used where the transcript start and end sites were marked as complete and the coding span was a multiple of three.

Likely benign variants were obtained through the ExAC browser. For simplicity, in this set we considered only single nucleotide polymorphisms. Variants were split into ‘common’ and ‘rare’ based on the maximum allele frequency present in any population with allele frequency of greater than 1 in 10,000 being defined as common and all others as rare. The March 16, 2017 release 3 of probability of loss of function intolerant predictions were also obtained from the ExAC browser.

When computing enrichments of ExAC variants, we wished to control for nucleotide content and distance from the 3’ss. This was particularly important as we noticed that T’s are overall less prone to variation, leading to an artificially strong signal at the −2 position. To accomplish this, we compared variant frequency at branchpoints with a UNA motif to variant frequency at UNA motifs not implicated as branchpoints. We then defined a variant enrichment as rate of variants for branchpoint UNAs divided by rate of variants for non-branchpoint UNAs. We computed the statistical significance at each position relative to the branchpoint using the two-sided Fisher Exact test available through Scipy. We computed these statistics for both allele frequency > 0.0001 and allele frequency <= 0.0001, branchpoints in pLI >= 0.9 genes and branchpoints in pLI < 0.0001 genes. We again used a fisher exact test to assess statistical significance between these cases at each nucleotide.

### Exon Type Annotations

The 2013 version 2 build of MISO exon skipping and intron retention event annotations were downloaded from the MISO wiki (https://miso.readthedocs.io/en/fastmiso/). We made no attempt to filter these annotations based on additional functional data.

### 3’ss Strength Quantification

We used the MaxEntScan package, as available at http://genes.mit.edu/burgelab/maxent/download/, to quantify the strength of 3’ss. An in-house script was developed to invoke the program cleanly in Python, but no functional changes were made.

## Data Access

Code to recreate all components of our study and final trained model weights are available at https://github.com/jpaggi/labranchor. A bed file of predicted branchpoints for Gencode v19 protein coding genes is available in **Supplemental Table 2**. A file of LaBranchoR scores for all positions 70 base pairs upstream of a 3’ss is in **Supplement Table 3**. *In silico* mutagenesis scores for the 70 base pairs upstream of all exons in Gencode v19 protein coding genes are in **Supplement Table 4**.

## Acknowledgements

We would like to thank Karthik Jagadeesh for helpful advice on obtaining and processing variant data and Peter Stenson and David Cooper for access to HGMD.

## Author Contributions

J.M.P. conceptualized the study, performed the analyses, and wrote the manuscript with G.B‥

---

**Supplementary Figure 1.**
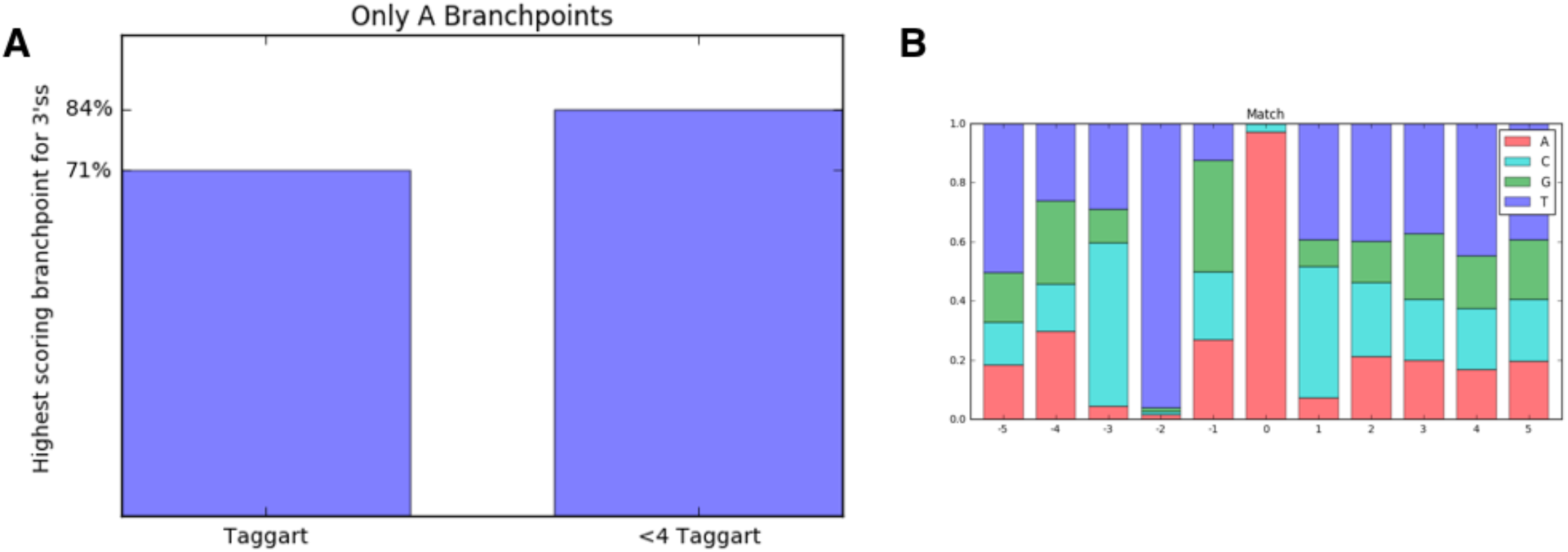
Overlap with Taggart branchpoints. (A) Fraction of predictions matching Taggart branchpoints with an A at the branchpoint position. (B) The overlap of Taggart and predicted branchpoints nearly invariantly have an A at the branchpoint position and U at the −2 U position.

**Supplementary Figure 2.**
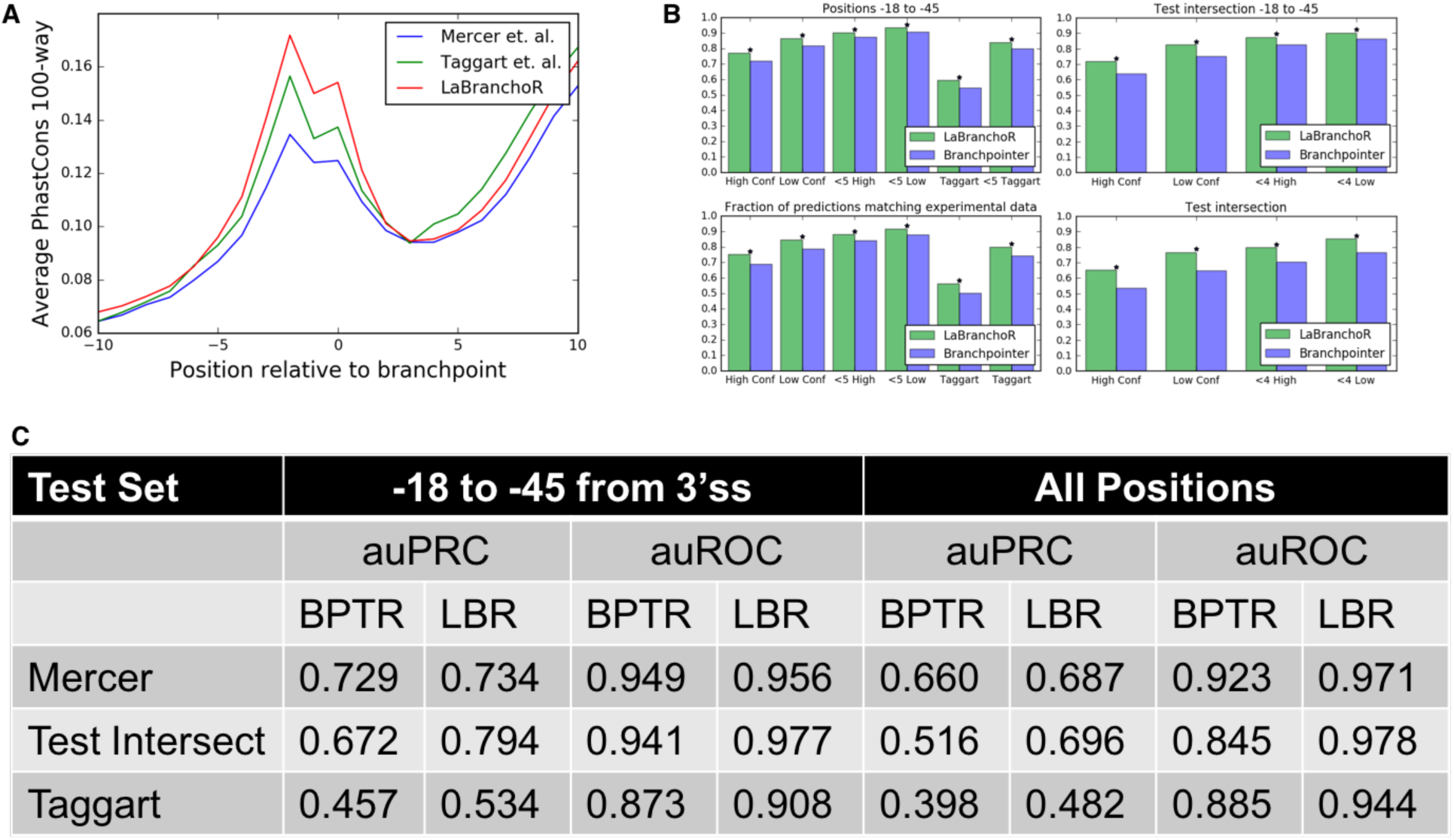
Evaluation of predictive performance and comparison to *branchpointer*. (A) PhastCons conservation signature for experimental and predicted branchpoints. (B) Fraction of predictions agreeing with Mercer branchpoints for positions −18 to −45 (top) and −5 to −60 (bottom) on our test set (left) and the intersection of our test set and the *branchpointer* test set (right). A * indicates a significant difference with p < 1e-4 by a one-sided Fisher Exact test. (C) Area under the precision-recall (auPRC) and receiver-operator (auROC) curve statistics for *branchpointer* (BPTR) and LaBranchoR (LBR). ‘Mercer’ refers to all Mercer high confidence sites. ‘Test Intersect’ is the intersection of our test set and the *branchpointer* test set. ‘Taggart’ is the set of Taggart et. al. 2017 branchpoints on chromosome 1-4.

**Supplementary Figure 3.**
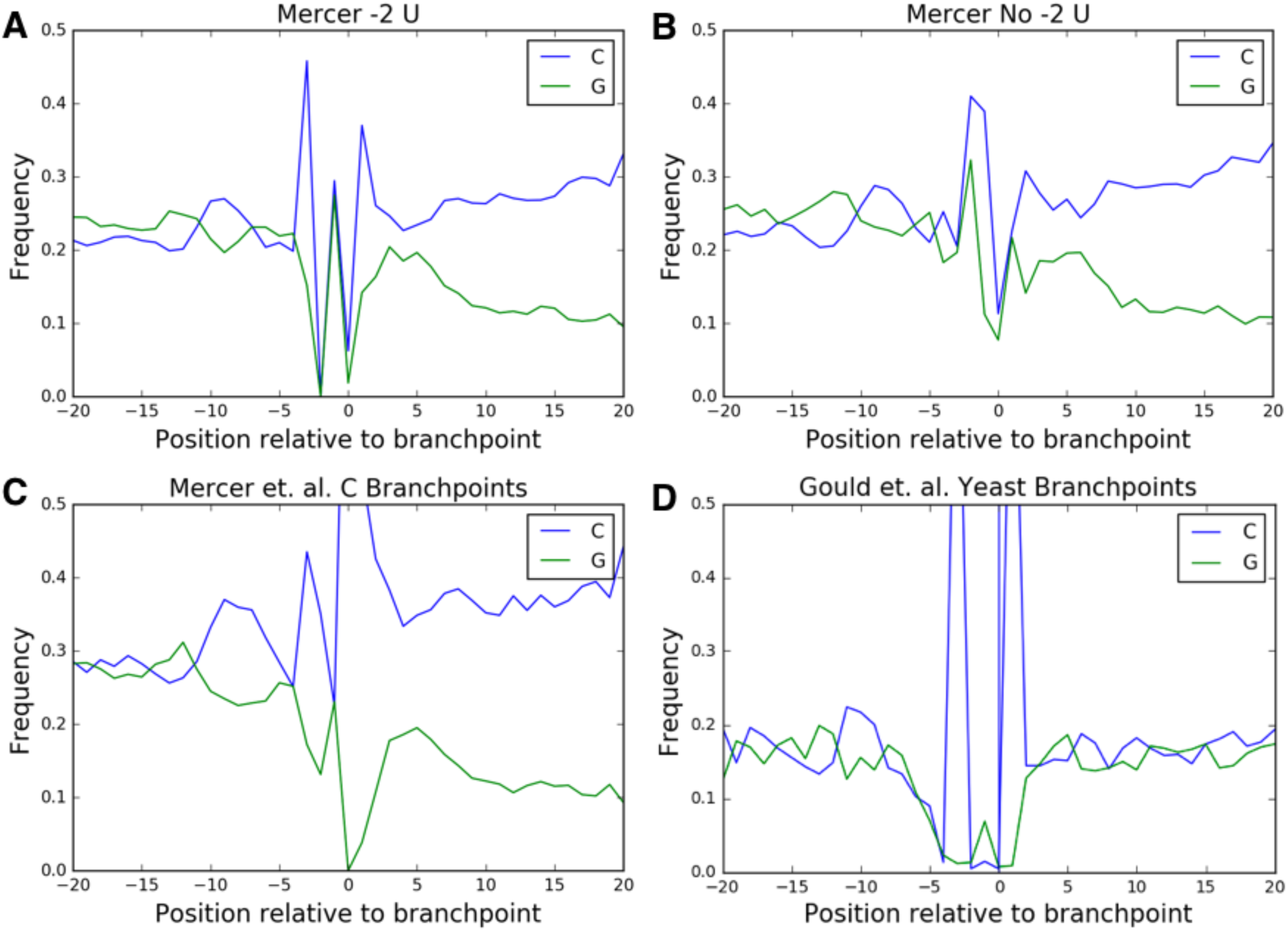
Experimentally determined branchpoints display a nucleotide content signature, consistent with extended base pairing with U2 snRNA. (A, B, C) Identical to Figure 3a-c, except for Mercer et. al. experimental data instead of predicitons. (D) G, C nucleotide content signature for a set of 718 experimentally determined budding yeast branchpoints.

**Supplementary Table 1.** Weak branchpoints play a role in alternative splicing. ‘Up SE’ correspond to branchpoints upstream of a conditionally skipped exon and ‘Down SE’ corresponds to branchpoints in introns downstream of conditionally skipped exons. ‘Retained’ refers to conditionally retained introns. Columns 2 and 3 refer to the fraction of branchpoints associated with each alternative splicing event. The enrichment and associated p-value (determined by a two-sided Fisher exact test) correspond to the ratio of columns 3 and 2.

| EVENT | A AT BP | C AT BP | ENRICH | P-VALUE |
| --- | --- | --- | --- | --- |
| UP SE | 0.1199 | 0.1452 | 1.210 | 1e-5 |
| DOWN SE | 0.1093 | 0.0990 | 0.906 | 0.062 |
| RETAINED | 0.0308 | 0.0727 | 2.356 | 8e-39 |

**Supplementary Table 1.** Weak branchpoints play a role in alternative splicing.
| EVENT | T AT -2 | NOT T AT -2 | ENRICH | P-VALUE |
| --- | --- | --- | --- | --- |
| UP SE | 0.1179 | 0.1311 | 1.1120 | 9e-13 |
| DOWN SE | 0.1111 | 0.1003 | 0.902 | 6e-10 |
| RETAINED | 0.0294 | 0.0406 | 1.380 | 5e-28 |

**Please see http://bejerano.stanford.edu/labranchor/ for the below tables**.

**Supplemental Table 2**. Genome-wide branchpoint predictions. A bed file of branchpoint predictions for all introns in protein coding genes in the Gencode v19 annotations. Each entry corresponds to the nucleotide with highest predicted probability of being a branchpoint for each 3’ss.

**Supplemental Table 3**. Genome-wide branchpoint probabilities. Similar to **Supplementary Table 2**, except contains branchpoint probabilities for all 70 nucleotides upstream of each 3’ss.

**Supplemental Table 4**. Genome-wide *in silico* mutagenesis scores. Mutagenesis scores for all possible single nucleotide polymorphisms in the 70 nucleotides upstream of all 3’ss in protein coding genes in the Gencode v19 annotations.

## References

1. Pascolo, E. & Séraphin, B. The branchpoint residue is recognized during commitment complex formation before being bulged out of the U2 snRNA-pre-mRNA duplex. Mol. Cell. Biol. 17, 3469–3476 (1997).

2. Hoskins, A. A. & Moore, M. J. The spliceosome: a flexible, reversible macromolecular machine. Trends Biochem. Sci. 37, 179–188 (2012).

3. Taggart, A. J., DeSimone, A. M., Shih, J. S., Filloux, M. E. & Fairbrother, W. G. Large-scale mapping of branchpoints in human pre-mRNA transcripts in vivo. Nat. Struct. Mol. Biol. 19, 719–721 (2012).

4. Taggart, A. J. et al. Large-scale analysis of branchpoint usage across species and cell lines. Genome Res. gr.202820.115 (2017).

5. Mercer, T. R. et al. Genome-wide discovery of human splicing branchpoints. Genome Res. gr.182899.114 (2015). doi:10.1101/gr.182899.114

6. Signal, B., Gloss, B. S., Dinger, M. E. & Mercer, T. R. Machine-learning annotation of human splicing branchpoints. bioRxiv 094003 (2016). doi:10.1101/094003

7. Alsafadi, S. et al. Cancer-associated *SF3B1* mutations affect alternative splicing by promoting alternative branchpoint usage. Nat. Commun. 7, ncomms10615 (2016).

8. Heintz, C. et al. Splicing factor 1 modulates dietary restriction and TORC1 pathway longevity in C. elegans. Nature 541, 102–106 (2017).

9. Hochreiter, S. & Schmidhuber, J. Long Short-Term Memory. Neural Comput. 9, 1735–1780 (1997).

10. Lipton, Z. C., Berkowitz, J. & Elkan, C. A Critical Review of Recurrent Neural Networks for Sequence Learning. ArXiv150600019 Cs (2015).

11. Bertram, K. et al. Cryo-EM structure of a human spliceosome activated for step 2 of splicing. Nature 542, 318–323 (2017).

12. Lim, K. H., Ferraris, L., Filloux, M. E., Raphael, B. J. & Fairbrother, W. G. Using positional distribution to identify splicing elements and predict pre-mRNA processing defects in human genes. Proc. Natl. Acad. Sci. 108, 11093–11098 (2011).

13. Yeo, G. & Burge, C. B. Maximum entropy modeling of short sequence motifs with applications to RNA splicing signals. J. Comput. Biol. J. Comput. Mol. Cell Biol. 11, 377–394 (2004).

14. Landrum, M. J. et al. ClinVar: public archive of interpretations of clinically relevant variants. Nucleic Acids Res. 44, D862–868 (2016).

15. Stenson, P. D. et al. The Human Gene Mutation Database: building a comprehensive mutation repository for clinical and molecular genetics, diagnostic testing and personalized genomic medicine. Hum. Genet. 133, 1–9 (2014).

16. Lek, M., Karczewski, Konrad J. & Minikel, Eric V. Analysis of protein-coding genetic variation in 60,706 humans. Nature 536, 285–291 (2016).

17. Xiong, H. Y. et al. The human splicing code reveals new insights into the genetic determinants of disease. Science 347, 1254806 (2015).

18. Shepard, P. J., Choi, E.-A., Busch, A. & Hertel, K. J. Efficient internal exon recognition depends on near equal contributions from the 3’ and 5’ splice sites. Nucleic Acids Res. 39, 8928–8937 (2011).

19. Gozani, O., Feld, R. & Reed, R. Evidence that sequence-independent binding of highly conserved U2 snRNP proteins upstream of the branch site is required for assembly of spliceosomal complex A. Genes Dev. 10, 233–243 (1996).

20. Yu, A. T., Ge, J. & Yu, Y.-T. Pseudouridines in spliceosomal snRNAs. Protein Cell 2, 712–725 (2011).

21. Gould, G. M. et al. Identification of new branch points and unconventional introns in Saccharomyces cerevisiae. RNA N. Y. N 22, 1522–1534 (2016).

22. Kierzek, E. et al. The contribution of pseudouridine to stabilities and structure of RNAs. Nucleic Acids Res. 42, 3492–3501 (2014).

